# Analysis-rcs-data: Open-source toolbox for the ingestion, time-alignment, and visualization of sense and stimulation data from the Medtronic Summit RC+S system

**DOI:** 10.1101/2021.06.07.447439

**Authors:** Kristin K Sellers, Ro’ee Gilron, Juan Anso, Kenneth H Louie, Prasad R Shirvalkar, Edward F Chang, Simon J Little, Philip A. Starr

**Author notes:** Shared first author (Kristin K Sellers, Ro’ee Gilron, Juan Anso). Shared senior author (Simon J Little, Philip A. Starr).

## Abstract

Closed-loop neurostimulation is a promising therapy being tested and clinically implemented in a growing number of neurological and psychiatric indications. This therapy is enabled by chronically implanted, bidirectional devices including the Medtronic Summit RC+S system. In order to successfully optimize therapy for patients implanted with these devices, analyses must be conducted offline on the recorded neural data, in order to inform optimal sense and stimulation parameters. The file format, volume, and complexity of raw data from these device necessitate conversion, parsing, and time reconstruction ahead of time-frequency analyses and modeling common to standard neuroscientific analyses. Here, we provide an open-source toolbox written in Matlab which takes raw files from the Summit RC+S and transforms these data into a standardized format amenable to conventional analyses. Furthermore, we provide a plotting tool which can aid in the visualization of multiple data streams and sense, stimulation, and therapy settings. Finally, we describe an analysis module which replicates RC+S on-board power computations, functionality which can accelerate biomarker discovery. This toolbox aims to accelerate the research and clinical advances made possible by longitudinal neural recordings and adaptive neurostimulation in people with neurological and psychiatric illnesses.

## Introduction

Bidirectional, chronically implanted, neural interfaces provide an unprecedented window into human neural activity during daily living and across a range of disease and symptom states. In addition, these devices can deliver therapeutic stimulation in response to real-time changes in neural activity features, driven by symptom biomarkers (Lo and Widge 2017; Velisar et al. 2019; Bouthour et al. 2019). Compared to traditional deep-brain stimulation (DBS) paradigms, this adaptive stimulation approach may provide more nuanced therapy, avoiding side effects and maximizing potential benefit (Huang et al. 2019; Herron et al. 2016; Swann et al. 2018; Velisar et al. 2019; Little et al. 2016). Furthermore, the neural sensing capability of bidirectional devices opens new possibilities for understanding disease mechanisms and functional brain networks (Swann et al. 2017). The Summit RC+S from Medtronic (Stanslaski et al. 2018), a device available under Investigational Device Exemption, is currently employed in the study of a wide range of clinical indications (Table 1). It is a leading example of advanced bidirectional neuromodulation technology that has heralded in a new era of longitudinal, high-volume brain sensing and neuromodulation in human patients. The advanced sense and stimulation capabilities of this device system provide great user flexibility, but also challenges for data handling. Data handling challenges include the need for critical software for reading, handling, processing, or analyzing RC+S data streams.

**Table 1:**
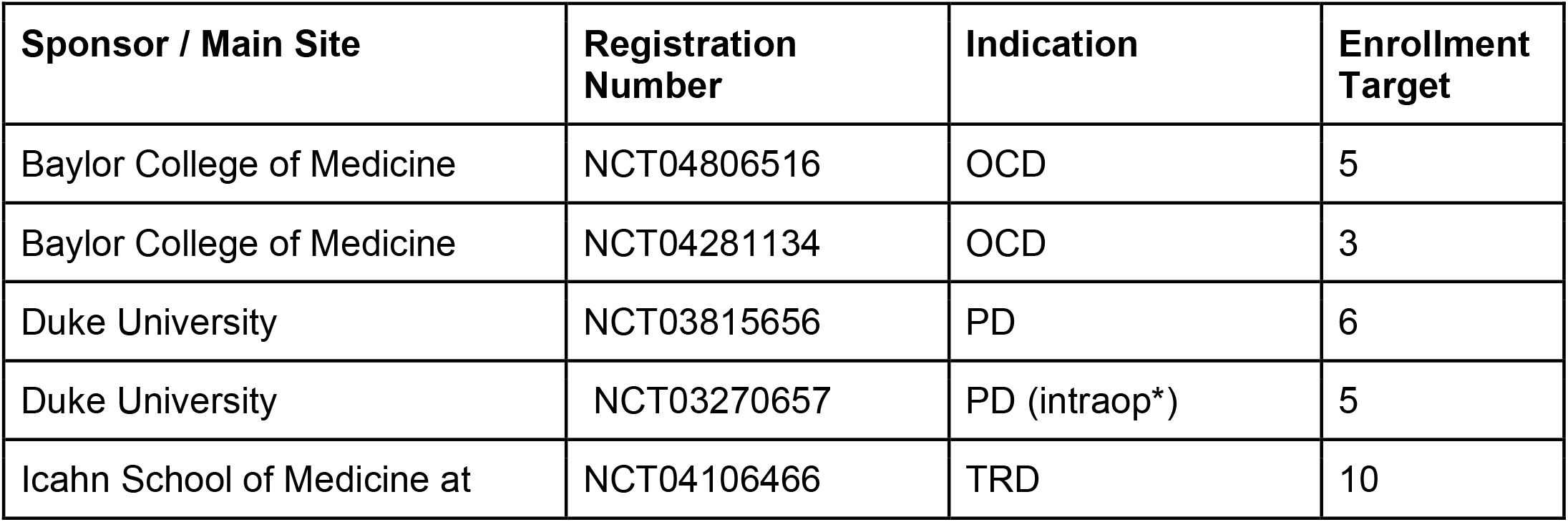
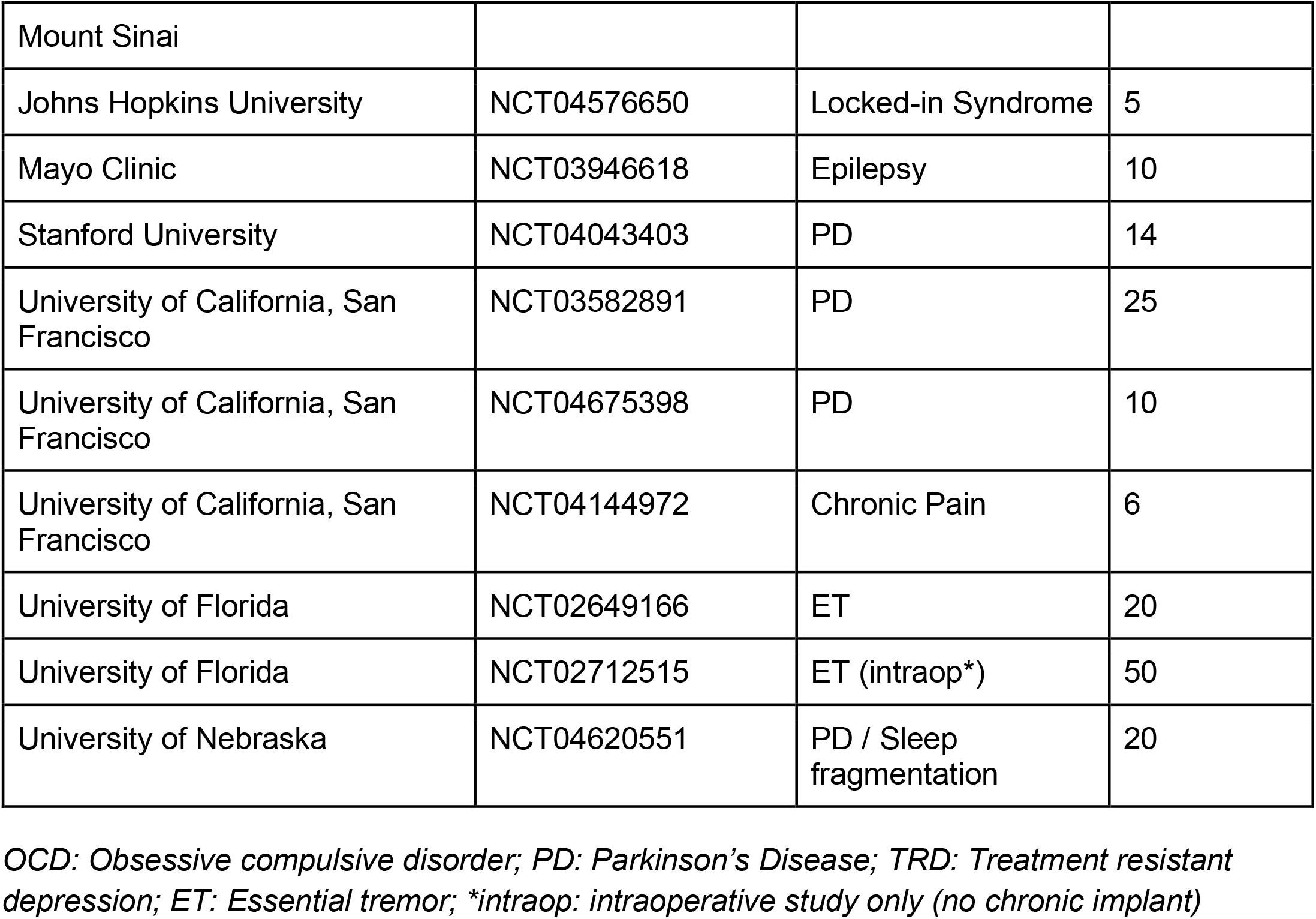
Clinical trials using the Medtronic Summit RC+S system

In order to prevent multiple individual research teams from needing to engineer piecemeal solutions specific to each use-case simply to access the data, we here provide a freely available, comprehensive software toolbox written in Matlab and tested on Mac and Windows (https://github.com/openmind-consortium/Analysis-rcs-data). We describe the implementation of this functionality in three parts, with example patient and benchtop data: (1) A data translation tool to ingest raw data from the Summit RC+S and transform those data into a user-friendly, human-readable, conventional analysis-ready format with data streams on a common time base, with consistent inter sample intervals; (2) A plotting tool that dynamically displays multiple raw data streams and associated metadata; and (3) An analysis module that mimics on-board power calculations conducted by the device and plugs in to the constructed human-readable data. Together, these tools can be used to support wide ranging analyses of RC+S data or modeling developed by the end-user.

### Medtronic Summit RC+S

The Summit RC+S system consists of two surface or depth leads that are implanted in the brain and a chest-located implantable neurostimulator (INS). The system is capable of sensing neural activity, performing on-board computations, and delivering open-loop or adaptive stimulation based on user-programmed parameters. The device can stream myriad metadata (device and battery status; sensing, stimulation, and adaptive configurations; enabled electrode contacts, etc.) in addition to user-defined selections of time series data (referred to here as ‘data streams’, including: time domain local field potentials, band-pass power, fast fourier transform [FFT], accelerometry, and adaptive algorithm settings; Table 2) to an external tablet.

**Table 2:** Summit RC+S Configurable Data Streams

|  |  |
| --- | --- |
| Time Domain | Continuous time domain data from up to 4 channels sampled at 250 or 500Hz, or from up to 2 channels sampled at 1000Hz. |
| Accelerometry | Continuous onboard 3-axis accelerometry data sampled at ~4 - 64Hz |
| FFT | Single-sided fast fourier transform derived by the on-device FFT engine according to user-defined FFT parameters |
| Power Domain | Continuous power data from the on-board FFT engine in configurable power bands. Up to 8 power domain channels can be streamed simultaneously. |
| Adaptive | Setting and stimulation state information from the adaptive detector. |

The richness and completeness of the data that is streamed also presents a number of challenges. The device employs User Datagram Protocol (UDP) to transmit packets of data from the implanted INS to an external tablet. However, this transmission protocol does not perform receipt verification, meaning that some data packets may be lost in transmission (e.g. if the patient walks out of range) and/or may be received out of order. Each of the packets contains a variable number of samples, and timing information is only present for the last sample in each packet (Figures 1A and C). These data packets are stored in 11 JSON files, such that 11 raw data files are present for each recording (Figure 1D). Packets are individually created, sent, and received for the different JSON files, meaning that packets across different data streams have different timing information, and missing packets across data streams may not align. The JSON files contain a combination of meta data and time series information with much of the metadata coded in hex or binary necessitating translation into human-readable values (Figures 1A and B). Lastly, information is needed from multiple JSON files simultaneously to provide users with information of interest (e.g. multiple JSON files are needed to recreate the labels of electrode contacts which were being used for stimulation and the parameters for stimulation) (Figures 1C and D). The quantity and variety of data from this device far surpass any previous bidirectional neuromodulation system, but this strength has also proven to be a notable barrier to implementation for research and clinical teams. The first and second parts of the presented toolbox seek to address this challenge by providing data parsing and time alignment across the data streams and streamlined data visualization.

**Figure 1:**
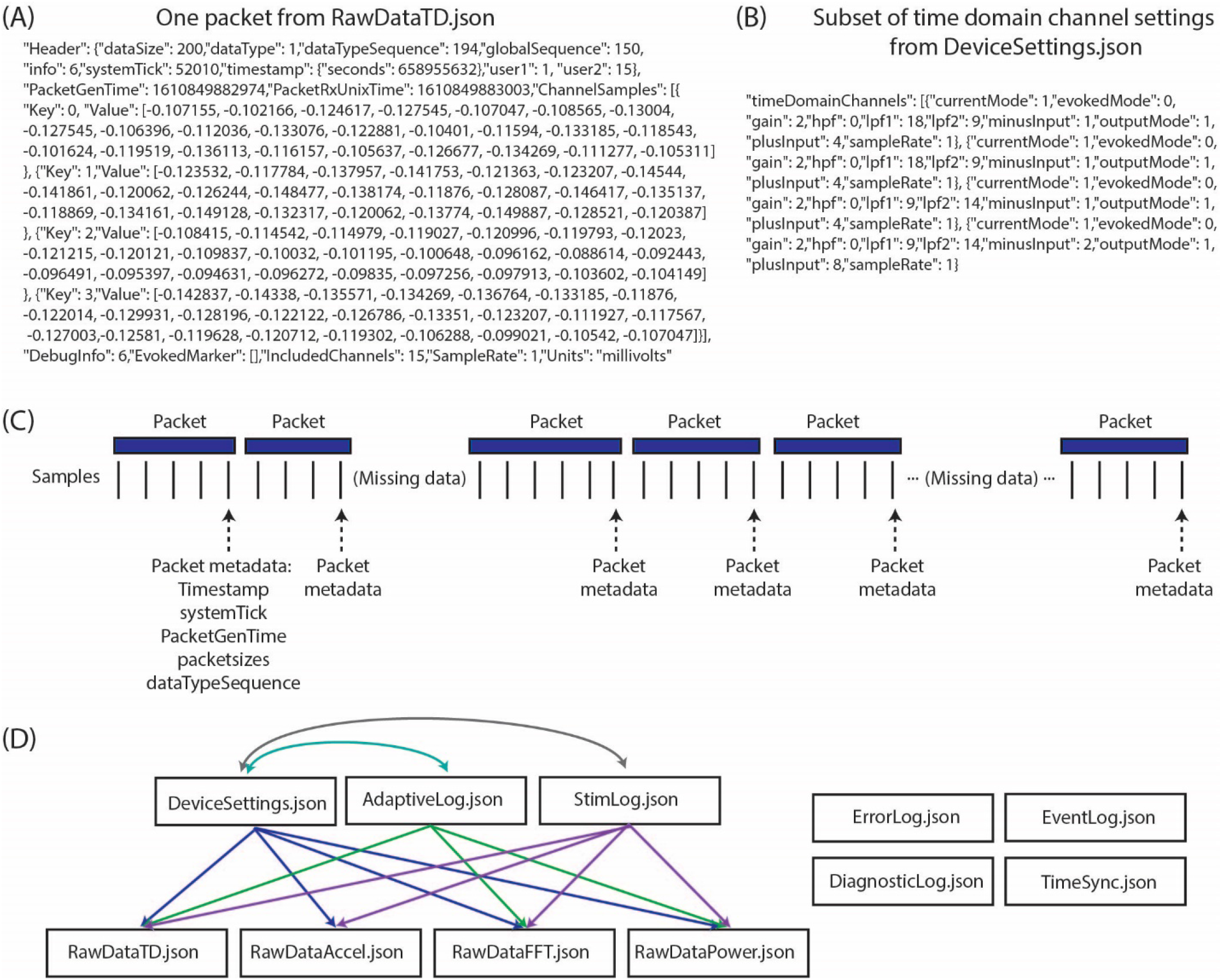
Summit RC+S raw data structure (A) One packet of data from RawDataTD.json; each packet contains one set of timing values and a variable number of time domain samples from each streamed channel; values such as SampleRate must be converted to interpretable values (e.g. Hz). (B) One section of values from DeviceSettings.json which provides information on time domain channel settings; mode, gain values, high pass and low pass filter settings, and contacts must be decoded to interpretable values. (C) Each time series stream transmits data from the INS in packets of variable sizes using UDP; receipt verification is not performed, so packets may not be received or may be received out of order. Each packet contains one value of timing information per variable, aligned to the last sample in the packet. Each data stream transmits packets separately, with non-aligned timing information. (D) The present toolbox is compatible with raw RC+S data which are acquired in 11 JSON files. This relationship diagram depicts that information from multiple files is required to interpret the recordings. For example, interpretation of RawDataTD.json may require all other JSON files which are connected to it via arrows. Colors are used to aid visualization.

A key mode of operation for the Summit RC+S uses an ‘embedded’ algorithm to control adaptive stimulation, which is also complex to implement. A typical workflow for programming of this mode includes identifying neural activity which is correlated or predictive of symptoms (i.e. a biomarker), programming the device to calculate the biomarker, and setting the device detector with threshold values such that when the biomarker moves between predefined states, stimulation delivery and/or stimulation parameters are adjusted. Specifically, the Summit RC+S includes on-board computational capability to calculate fast fourier transform (FFT), band-pass power, and execute linear discriminant analysis (LDA) to control the administration of adaptive stimulation. Effectively programming the device and managing patients using adaptive stimulation can be challenging because the biomarker characteristics (e.g. frequency band limits, dynamic range) must be known, and parameters of the on-board computation of the FFT and power (e.g. interval, size, hann window) can change values going into the LDA. Exhaustively testing these parameters in patients is time consuming and not feasible. Therefore, the third part of our toolbox is to provide a power calculation module which allows for off-device power computation using streamed time domain data. This tool can be set to use the same parameters as the Summit RC+S, allowing for the optimization of settings to increase detector performance without creating undue burden on the patient. A key feature differentiating the power computations in our toolbox from standard offline power calculations is that the magnitude, update rate, and range of power values will be comparable to those calculated by the device; these values can directly inform optimal programming of adaptive stimulation.

## Method and Results

### Part 1: Data Parsing and Time Alignment

Conventional neurophysiological analyses are greatly simplified by the use of a standardized timebase across data streams and a consistent sampling rate (i.e. inter-sample interval). This facilitates time-frequency decomposition and supports downstream modeling of disease biomarkers, analysis of stimulation impact, and parameter selection for adaptive stimulation. Such standardized data formatting includes data in matrix form, with samples in rows, data features in columns (or vice versa), and a timestamp assigned to each row. A key computational step for RC+S data is the derivation of the precise time assigned to each row, which we will refer to as DerivedTime. DerivedTime should be in unix time (a standardized time format for describing a point in time; the number of elapsed seconds from 1 January 1970 in UTC, with a method to account for different time zones), to allow for synchronization with external data streams, symptom reports, or tasks. Furthermore, we ideally would like all separate datastreams to be on the same timebase, aligned to common DerivedTime timestamps (such that we can analyze multiple data streams recorded simultaneously - for example correlating time and power domain data with patient movement detected via the accelerometer). Below, we describe our implemented approach to navigate the specialized native format of RC+S data to achieve this desired, standardized output format (Figure 2).

**Figure 2:**
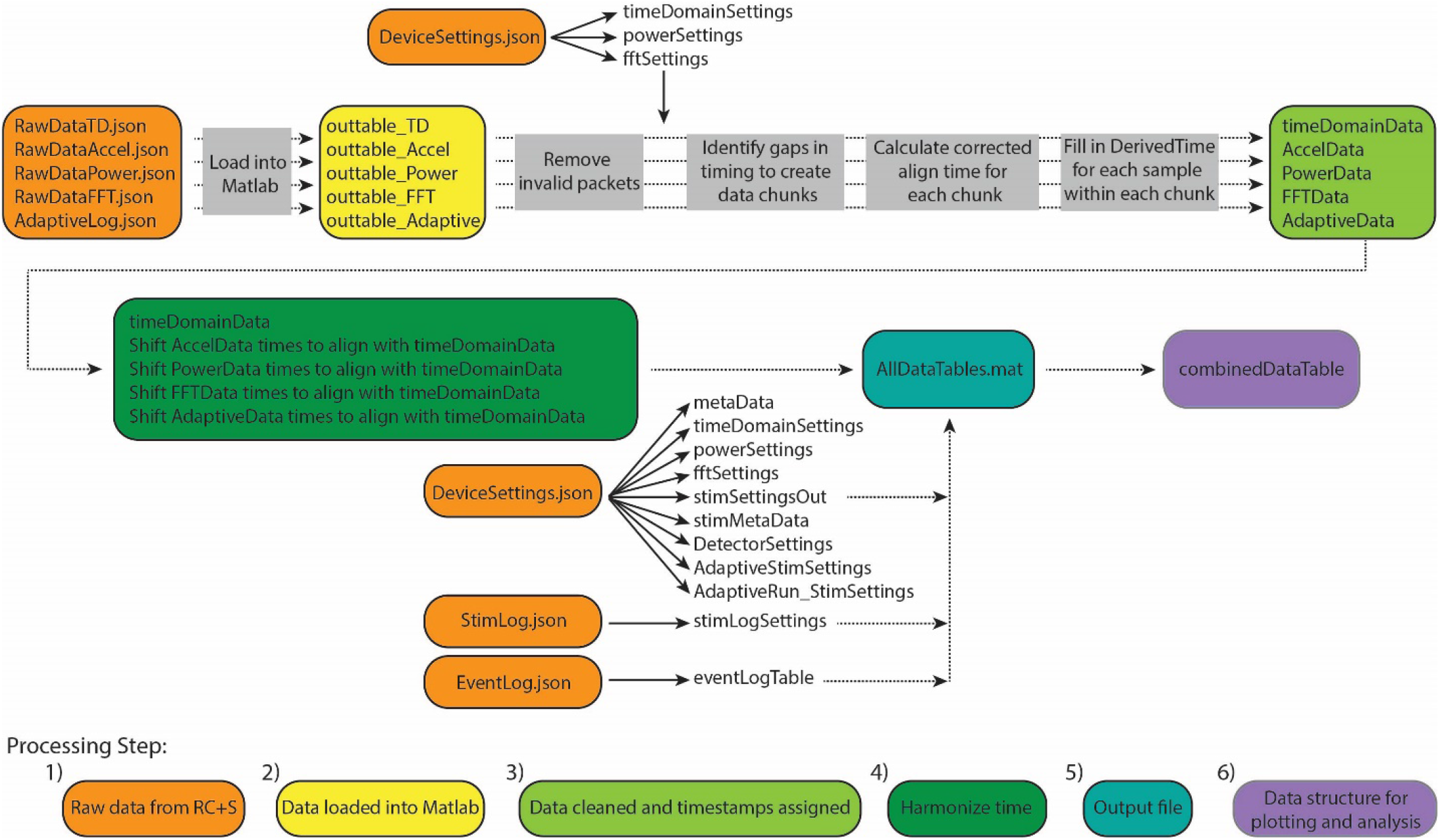
Overview of Summit RC+S data parsing and time alignment. Raw JSON files (orange) are loaded into Matlab (yellow). For each time series data stream, packets with invalid data are removed and timing variables are used to calculate DerivedTime for each sample (light green). Samples in each data stream are aligned to DerivedTime for time domain data, which has the highest sampling rate (dark green). These data tables are saved in a .mat file (using a combination of tables and sparse matrices) along with tables containing settings information and metadata (blue). Finally, combinedDataTable is created which can be used for plotting and user-specific analyses (purple).

The result of this approach is to provide a table (combinedDataTable) containing time series data from all data streams with a calculated DerivedTime value for each sample, and tables with relevant metadata and settings which can be applied to select periods of interest in combinedDataTable. DerivedTime is inclusive of the beginning of the earliest starting data stream to the end of the latest finishing data stream, in steps of 1/Fs of the time domain data stream (Fs = 250, 500, or 1000Hz). CombinedDataTable is filled with data from all datastreams; if there is not a sample for a given time step, the entry is filled with a NaN. Thus, this neuroscience-analysis-ready table can be quite large to store on disk (leading to prohibitively long read/write times for long recordings). Therefore, there are two main functions to execute to achieve the desired final data table: ProcessRCS.m followed by createCombinedTable.m (Table 3). In the following sections, we describe the rationale behind the implementation of these functions.

**Table 3:** Description of functions for creating CombinedDataTable

| Function | Inputs | Outputs |
| --- | --- | --- |
| ProcessRCS.m | (1) Path to folder containing raw JSON files<br>(2) [Optional] processFlag to indicate saving/read/overwrite selection<br>(3) [Optional] Alternate method for handling short gaps in data, for advanced users (more information below) | For each data stream: sparse matrix with numerical data, cell array with column labels for sparse matrix, table with non-numerical data; tables with metadata and settings |
| createCombinedTable.m | All required variables available from AllDataTables.mat or output of ProcessRCS.m<br>(1) Cell array of data streams to be included<br>(2) unifiedDerivedTimes<br>(3) metaData | combinedDataTable |

#### Steps 1 and 2: Raw data from RC+S loaded into Matlab

Large raw data are loaded from JSON files into Matlab using the turtle_json toolbox (https://github.com/JimHokanson/turtle_json, included in our toolbox repository), which can parse large files rapidly. In cases where JSON files are malformed (typically with closing brackets omitted), fixes are attempted to read these data. Each data stream is read independently, and empty or faulty raw data files will result in continuation of processing omitting that data stream.

#### Step 3: Data cleaned and timestamps aligned

We continue processing of each data stream independently. There are multiple time and counting related variables present for each packet of data (Table 4). We identify and remove packets with meta-data that is faulty or which indicate samples will be hard to place in a continuous stream (e.g. packets with timestamp that is more than 24 hours away from median timestamp; packets with negative PacketGenTime; packets where PacketGenTime goes backwards in time more than 500ms; packets where elapsed PacketGenTime disagrees with elapsed timestamp by more than 2 seconds).

**Table 4:** Time and count variables associated with each packet of data from the RC+S system

| Variable | Value meaning | Why insufficient for DerivedTime |
| --- | --- | --- |
| timestamp | Elapsed number of seconds since March 1, 2000, in units of seconds. Implemented in INS firmware | Highest resolution is 1 sec |
| systemTick | Running counter, in units of $1e4$ seconds; rolls over every $2^{16}$ values ($\sim 6.5535$ sec). Implemented in INS hardware | Rolls over every $2^{16}$ values. |
| PacketGenTime | API estimate of when the packet was created on the INS. Unix time with resolution to millisecond | The difference between adjacent PacketGenTime values does not always equal the expected amount of elapsed time. Aligning by PacketGenTime would result in varying inter-sample intervals |
| PacketRxUnixTime | Unix time when computer received packet | Highly inaccurate after packet drops |
| dataTypeSequence | Packet sequence number. Rolls over at 255; does not reset upon start of streaming | Counter, does not provide time |

Upon inspection of empirical patient and benchtop (Powell et al. 2021; Stanslaski et al. 2012) data sets, we found that none of the time related variables associated with each packet of data could independently serve as DerivedTime. Table 4 describes why each variable cannot be used for DerivedTime. In the case of PacketGenTime, the difference between PacketGenTime of adjacent packets, when no packets were dropped, does not equal the expected amount of elapsed time (as calculated using the number of samples in the packet and the sampling rate); the amount of this offset varies between packets. This presents a serious problem – in cases of missing time, we would lose the stereotyped 1/Fs duration between samples, which would introduce artifacts in time-frequency decomposition. In cases of overlap, there is no way to account for having more than one value sampled at the same time.

We next sought to use timestamp and systemTick in concert to create DerivedTime, and then convert to unix time using one value of PacketGenTime. However, we observed from empirical data (both recorded from an implanted patient device and using a benchtop test system) that one unit of timestamp (1 second) did not always equal 10,000 units of systemTick. The consequence of this was offset between systemTick and timestamp that accumulated over the course of a recording (multiple seconds error by the end of a 10-hour recording). While using these values may be acceptable for short recordings, we chose to move away from this implementation because one of the strengths of the RC+S system is the ability to stream data for long periods of time. Thus, rather than use any one of these time variables independently, we rely on information provided by all of them to create DerivedTime.

Our implemented solution for creating DerivedTime (Figure 3) depends on first identifying continuous “chunks” of data; defined as a continuous series of packets of data sampled without packet loss. Although there is indeterminacy in the timing of individual data packets, the INS device samples continuously at a fixed sampling interval and therefore, within a chunk of concatenated packets, the data sampling is continuous and regular. Our approach aims to align the beginning of continuous chunks of data to unix time and then use the sampling rate to determine the DerivedTime for each individual sample. This process relies on the assumption that only full packets of data are missing, but there are no individual samples missing between packets. First, we chunk the data - identified by looking at the adjacent values of dataTypeSequence, timestamp, and systemTick as a function of sampling rate and number of samples per packet. Breaks between chunks can occur because packets were removed during data cleaning, because there were dropped packets (never acquired), or because streaming was stopped but the recording was continued. Changes in time domain sampling rate will also result in a new chunk. There are two categories of chunks, short-gap and long-gap. Short-gap chunks follow a gap shorter than 6 seconds, as determined by timestamp (indicating there was not a full cycle of systemTick); long-gap chunks follow a gap greater than or equal to 6 seconds (indicating there may have been a full cycle of systemTick). There are two options for how to handle short-gap chunks and only one method for handling long-gap chunks.

**Figure 3:**
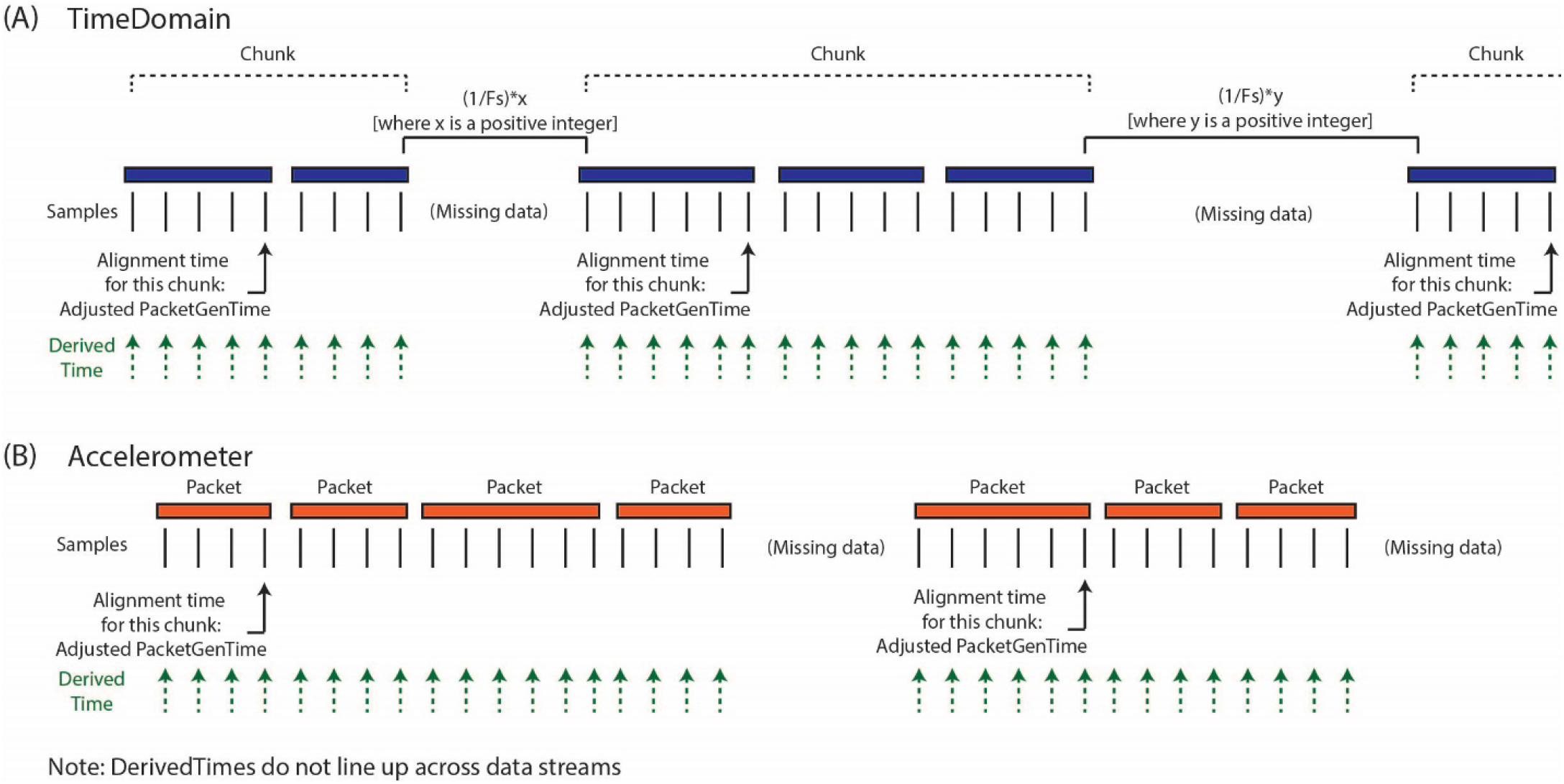
Calculation of DerivedTimes for each data stream (A) The default method for calculating DerivedTimes for short-gap chunks and the only method for long-gap chunks is to align the beginning of continuous chunks of data to Unix time using the adjusted PacketGenTime from the first packet in the chunk, and then using the sampling rate to determine the DerivedTime for each sample. Each DerivedTime is shifted to the nearest multiple of 1/Fs after chunk one in order to preserve consistent intersample spacing. (B) DerivedTime is calculated separately for each time series data stream, as each data stream has packets that are sent independently.

For all chunks, we need to align the beginning of the chunk to a Unix time. The first chunk in a recording is aligned using the PacketGenTime of the first packet in the chunk. The default option for handling short-gap chunks is the use of the same approach used for long-gap chunks: we look across all the packets in the chunk and calculate the average offset between each PacketGenTime and the amount of time that is expected to have elapsed (calculated based on sampling rate and number of samples in the packet). We then apply this offset to the PacketGenTime corresponding to the first packet of the chunk, creating the Adjusted PacketGenTime. We can now calculate a time for each sample in the chunk, as a function of the sampling rate. The alternative option for short chunks is to use adjacent values of systemTick to calculate the elapsed time across a gap (systemTick from the last packet of the previous chunk and systemTick of the first packet of the next chunk). This is possible because we have stayed within one full cycle of systemTick values. This approach should only be used when users have verified that their systemTick clock is quite accurate (otherwise error can accumulate over the course of the recording). Whichever process is selected is repeated separately for each chunk.

Lastly, we shift the calculated DerivedTime values slightly for chunks two onwards, in order to match the time base of the sampling of the first chunk of data and preserve inter-sample spacing of 1/Fs. Any missing values are filled with NaNs. Again, the above processing is conducted separately for each data stream, as each of these streams have separate systemTick, timestamp, and PacketGenTime values reported per packet. Harmonization of DerivedTime across data streams is conducted later.

#### Step 4: Harmonize time

As described above, the optimal format for neuroscience-analysis-ready data is matrix form, with samples in rows, data features in columns, and a timestamp assigned to each row.

After creating DerivedTime separately for each time series data stream, we must ‘harmonize’ these times across data streams. By this, we mean samples in each data stream are aligned to the nearest value of DerivedTime from time domain data, which has the highest sampling rate (Figure 4). In some cases, data streams may extend before or after time domain data -- in these instances, we add values to DerivedTime in steps of 1/Fs (time domain Fs) to accommodate all samples.

**Figure 4:**
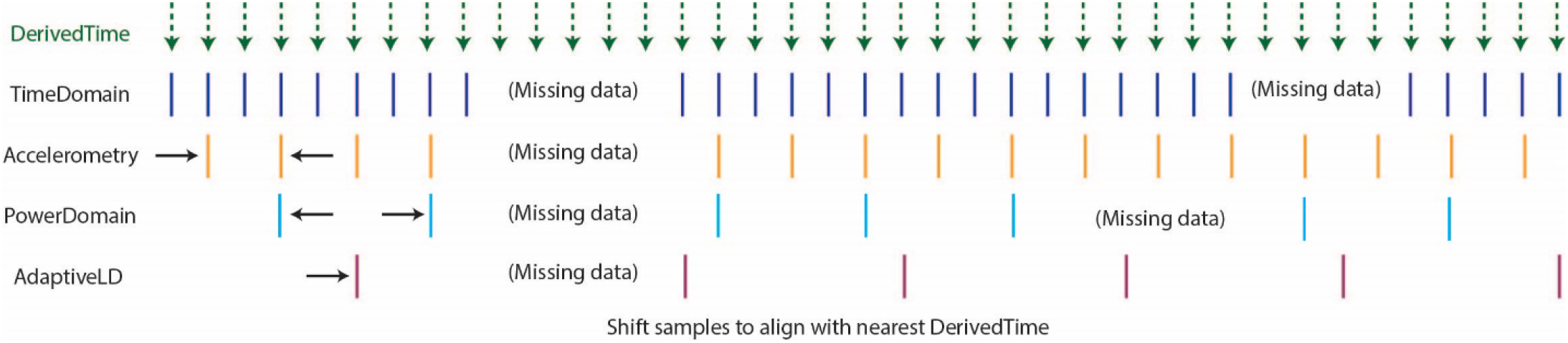
Harmonization of time across data streams to achieve one common DerivedTime timebase. DerivedTime from the time domain is taken as the common time base, as the time domain data have the highest sampling rate. Samples from other data streams are shifted in time slightly to align with the nearest time domain DerivedTime.

#### Step 5: Output file

If the user selects to save the output of ProcessRCS.m to disc, AllDataTables.mat is created and stored. This file contains a number of variables, which separately store data from each datastream and tables with metadata and settings. For each time series, numerical data are stored in a sparse matrix, non-numerical data are stored in a table, and a cell array contains the column headings of the sparse matrix. The purpose of saving these data broken into different tables and matrices is to minimize file size (as the final desired combinedDataTable contains a large number of NaNs and can be quite large).

#### Step 6: Data structure for plotting and analysis

Outputs from ProcessRCS.m (or variables loaded from AllDataTables.mat) can be used to create combinedDataTable using the script createCombinedTable.m. Whenever a data stream lacks a value for a particular DerivedTime, that entry in the table is filled with a NaN. The table does not contain any columns which are entirely filled with NaNs. The CombinedDataTable variable represents the final data structure for plotting and analysis. All time series data for a given session of RC+S streaming can be contained within this table. The use of Unix time facilitates the synchronization of neural data with external tasks, symptom reports, or across multiple implanted devices. For example, some patients are implanted with two RC+S devices (one in the right hemisphere, one in the left hemisphere) which can be streamed simultaneously. In Figure 5, we plot the accelerometry channels from bilateral devices in a single patient after each dataset was independently processed using ProcessRCS.m and combineCombinedTable.m. The movement signals are very closely aligned in time at the beginning and end of an overnight recording, providing an example of validation of the processing algorithm.

**Figure 5:**
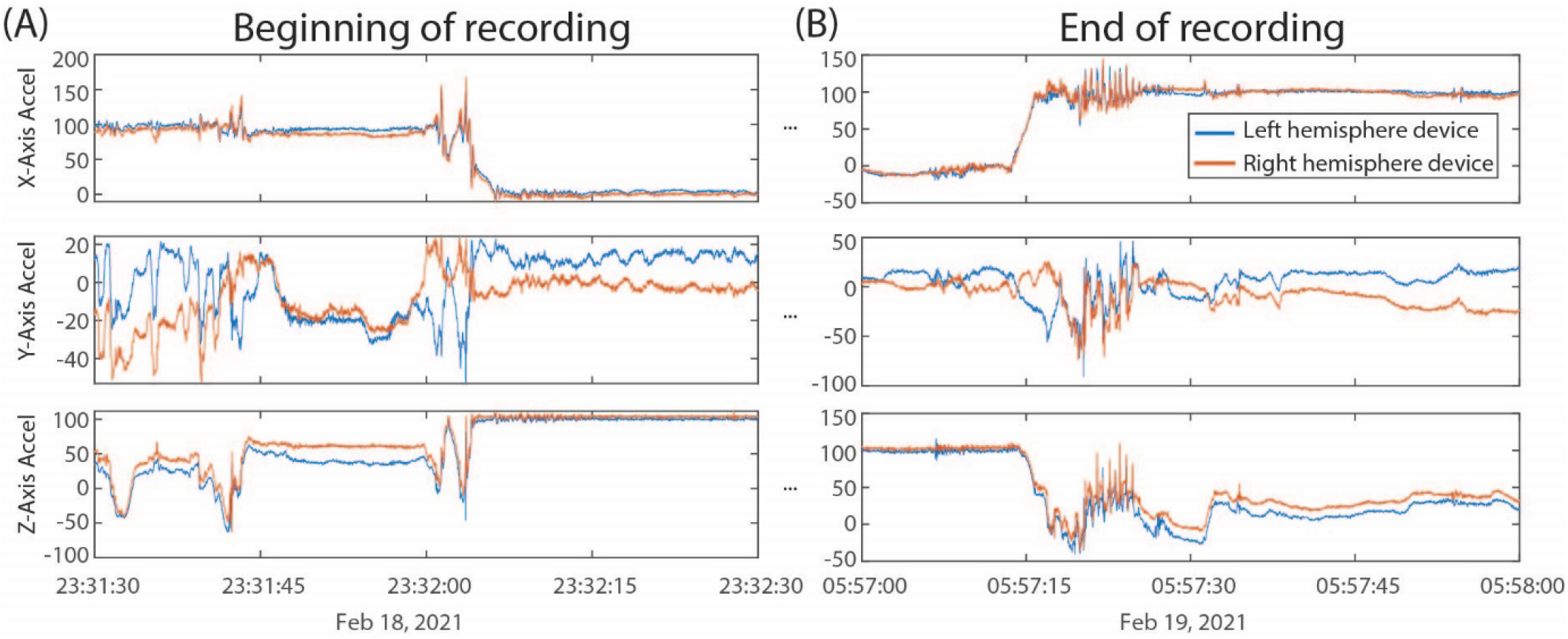
(A) Accelerometry channels from the beginning of an overnight recording from two RC+S devices implanted in the same patient. Detected movement serves as a way of confirming the parsing algorithm, which was applied separately to data from each device, is faithfully recreating time across the recording, without any accumulated offset. (B) Accelerometry channels from the end of the same ∼6.5 hour recording as in (A). No accumulated drift is visible between the datastreams across the devices.

### Part 2: Data Plotting and Visualization

Analysis of local field potential neural data often consists of several key steps: preprocessing, artifact removal, and spectral analysis. Performing these steps with the Summit RC+S data presents special challenges for a few key reasons: First, small gaps in the data introduce transient artifacts in spectral analysis. Second, RC+S data contains several data streams that are not commonly used in other processing and plotting pipelines (e.g. power time series, adaptive detector). Third, all data streams use different sampling rates. Fourth, data collected at home over hours and days (Gilron et al. 2021) result in multiple recording sessions; some analyses require loading multiple sessions and creating one cohesive structure. Finally, some data streams are usefully plotted together, such as the adaptive detector and associated thresholds.

In order to address these challenges, we have created a Matlab plotting tool to aid in rapidly plotting and analyzing RC+S data directly from the JSON files. Our plotting tool incorporates the functional steps described above to create a cohesive, unified time across RC+S data streams and provides the user the ability to easily plot all data types (Figure 6A). Unlike commonly available spectral tools (Fieldtrip, EEGLAB) which assume data are continuous, this tool will perform “gap aware” analysis of the data in the frequency and spectral domain. Data are plotted from multiple data streams with different sampling rates such that alignment is preserved, utilizing the common time base calculated in the first part of the toolbox, described above. Furthermore, we provide an easily executed mechanism to combine and analyze data from multiple sessions (e.g. throughout an entire day of streaming), as well as functions to save and aggregate power spectral density data for downstream analysis (Figure 6B). The plotting tools takes advantage of all meta-data parsing and combines this information in the display of plotting results. For example a call to plot a time domain channel will include information of the sense channels and filtering settings (Figure 6C, top), and a call to plot current will include information about stimulation channels, stimulation settings, and if changes occurred within the session (Figure 6C, second from bottom).

**Figure 6:**
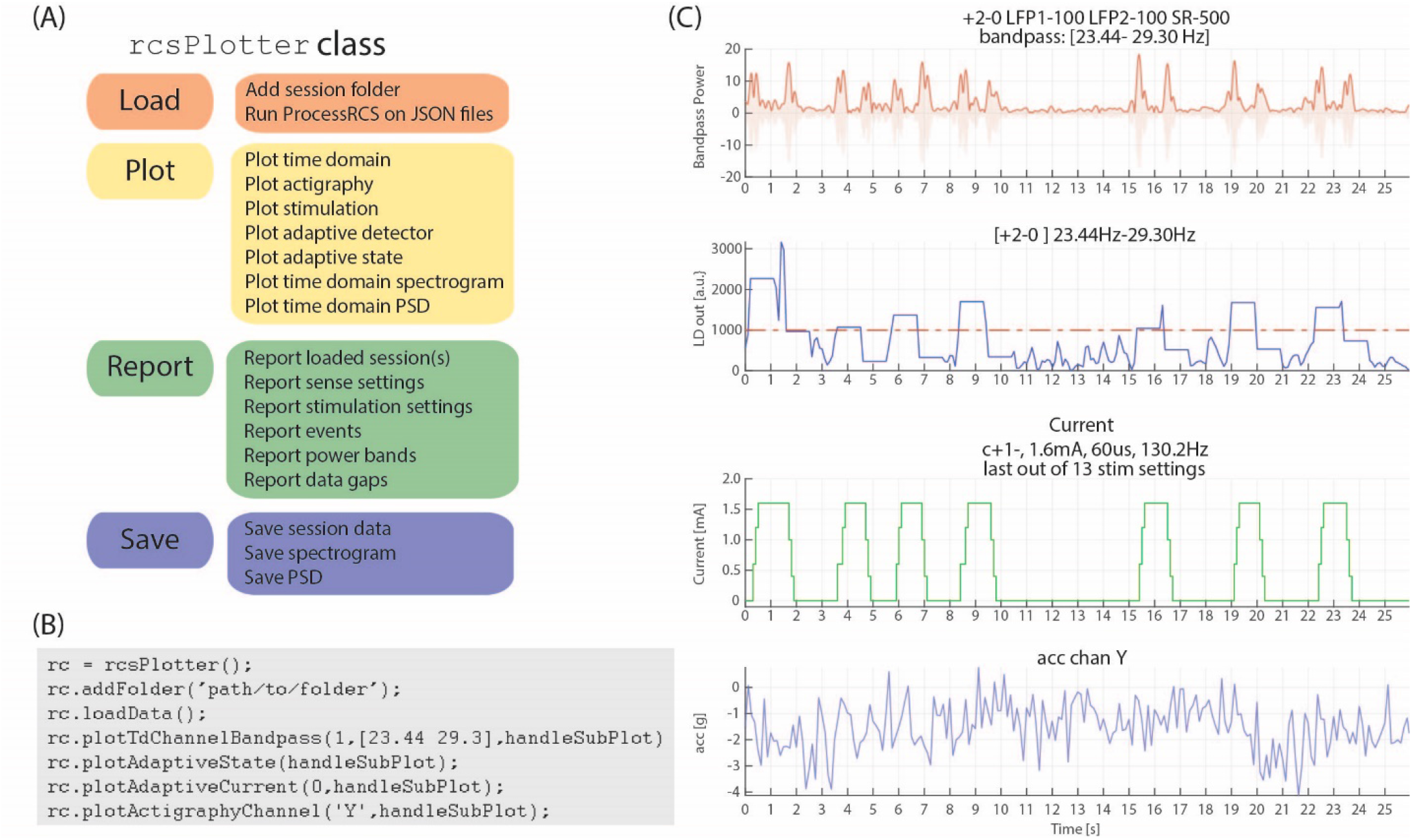
rcsPlotter overview and example (A) Main functions used in the ‘rcsPlotter’ class. These functions are used for loading data which are processed through ProcessRCS.m, plotting all RC+S data streams, reporting values across recordings (such as stim state and event markers), and saving for downstream analyses. (B) Example function call for the ‘rcsPlotter’. This shows the simplicity of loading data from an embedded adaptive DBS session and plotting the results. Plots from function call show in C. Each stream has its own dedicated plotting command that will pull in meta data and display it in the subplot title. Adding additional folders (for example, from the same day) only requires one call (and will plot all streams together). There is a “plot” method for each data stream. A list of available methods is available in the function help section. (C) This output from the ‘rcsPlotter’ class includes meta-data parameters pulled from multiple JSON files to populate graph titles. Top plot - bandpass time domain data used for embedded detector, sense channels and filter settings indicated. Second from top - output from embedded linear detector output (threshold shown as red dashed line). Third from top - stimulation current and current parameters. Bottom - actigraphy.

Finally, reporting functions exist to visualize gaps that exist in the data and report event markers (written to the raw JSON file by the API) that the experimenter may have programmed. Typically, these include task timing or patient symptom reports. Figure 6 provides a schematic of the analyses this tool can perform for data visualization as well as an example call demonstrating the simplicity of use to plot rich data stream visualizations.

### Part 3: Power calculation analysis module

The Summit RC+S can be programmed to deliver adaptive stimulation controlled by user-programmed power features and detector settings employing linear discriminant analysis. Biomarker discovery and programming of adaptive stimulation are greatly aided by being able to compute inferred embedded power domain outputs from the recorded time domain data off the device. This avoids the need for new data sets to be collected after any changes in device sense settings. Here, we describe an analysis module to calculate off-device power equivalent to the on-device power values using the streamed time domain neural data. This provides an estimate of power that is comparable to the power the device calculates internally and allows the user to calculate different frequency bands and with the option to modify FFT parameters (size, interval, Hann window %). Figure 7 provides an overview of the key computation steps in this module.

**Figure 7:**
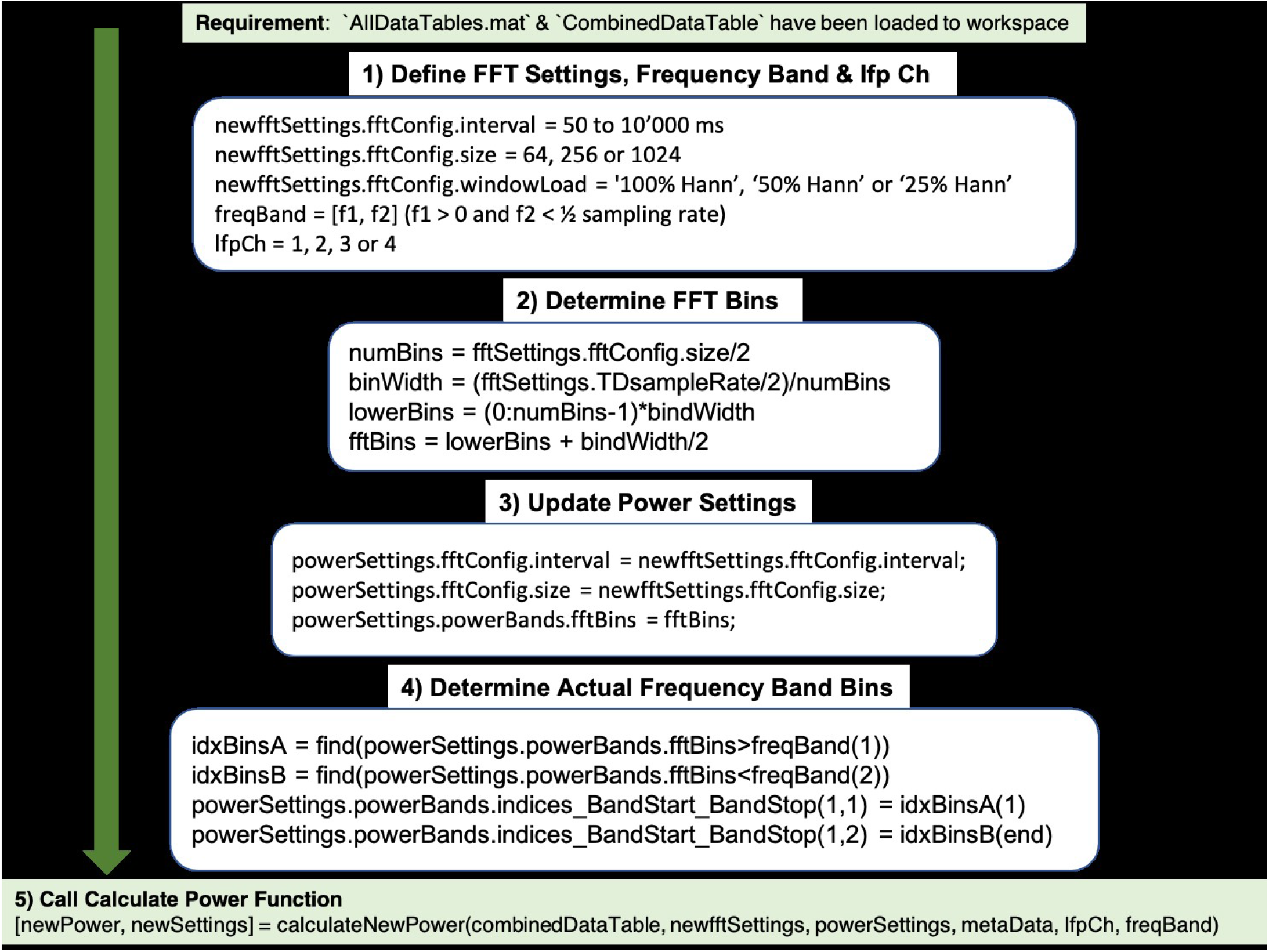
Use of the function calculateNewPower.m to calculate a new power domain time series based on user-defined FFT settings, frequency band, and time domain channel. The steps required before invoking the function include: (1) Define FFT settings, frequency band, and time domain channel. (2) Calculate FFT bins. (3) Define Power Settings using the FFT settings and derived FFT bins. (4) Determine FFT bins within frequency band. (5) Run function calculateNewPower.m passing all required parameters.

For the off-device power calculation, time domain signal s(n) is extracted from combinedDataTable, offset voltage is removed, and raw millivolt values are transformed to internal device units using the following equation which accounts for amplifier calibration (Eq 1; Table 5):

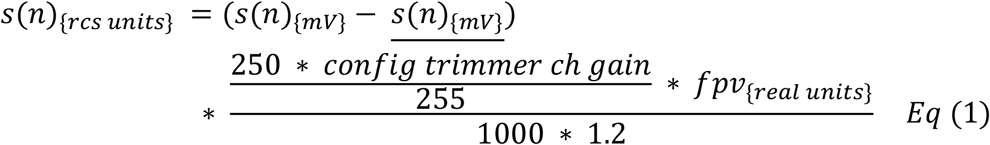

**Table 5:**
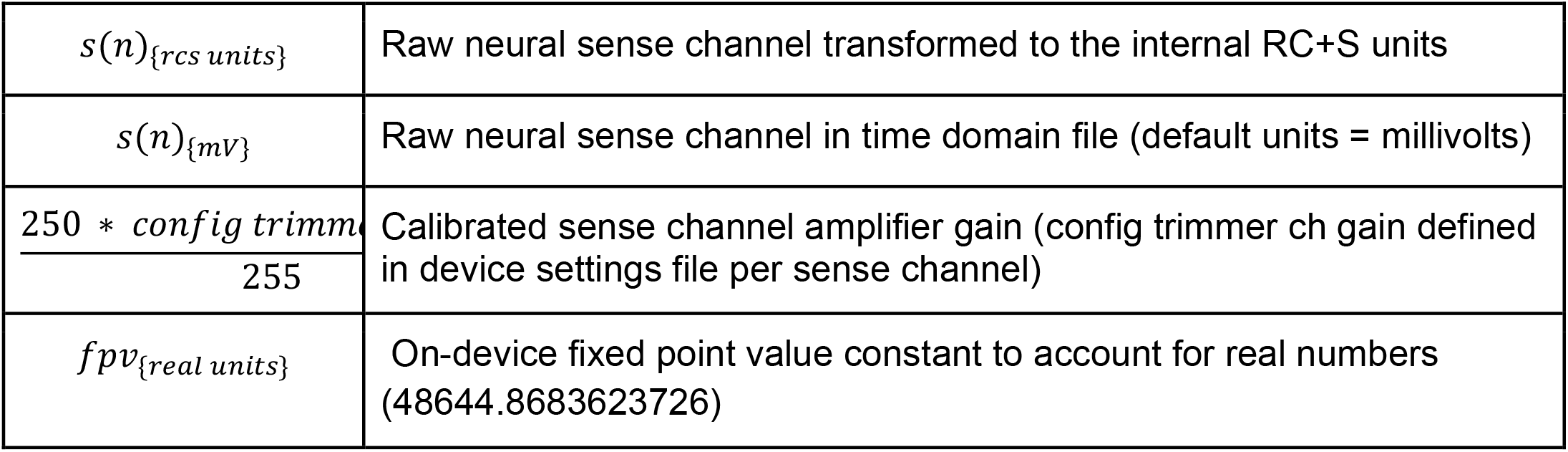
Variables and constants to transform RC+S signal back to internal on-device units.

|  |  |
| --- | --- |
| $s(n)_{\{rcs\ units\}}$ | Raw neural sense channel transformed to the internal RC+S units |
| $s(n)_{\{mV\}}$ | Raw neural sense channel in time domain file (default units = millivolts) |
| $\frac{250 * config\ trimm}{255}$ | Calibrated sense channel amplifier gain (config trimmer ch gain defined in device settings file per sense channel) |
| $fpv_{\{real\ units\}}$ | On-device fixed point value constant to account for real numbers (48644.8683623726) |

Then, the overlap of a running Hann window is calculated as a function of sampling rate, FFT interval, and FFT size. The overlap formula is given in (Eq 2):

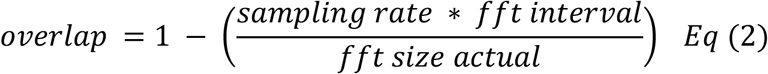

For the overlap calculation the device uses an actual number of FFT points of 62, 250, or 1000 for FFT sizes of 64, 256, or 1024, respectively. The RC+S offers three Hann windows (window load) settings, 25%, 50%, and 100%. The 100% Hann window is the default Hann window, defined by:

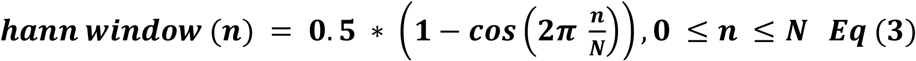

with a window length L = N + 1. In the off-device power calculation the user chooses one of the three Hann window settings (Figure 8).

**Figure 8:**
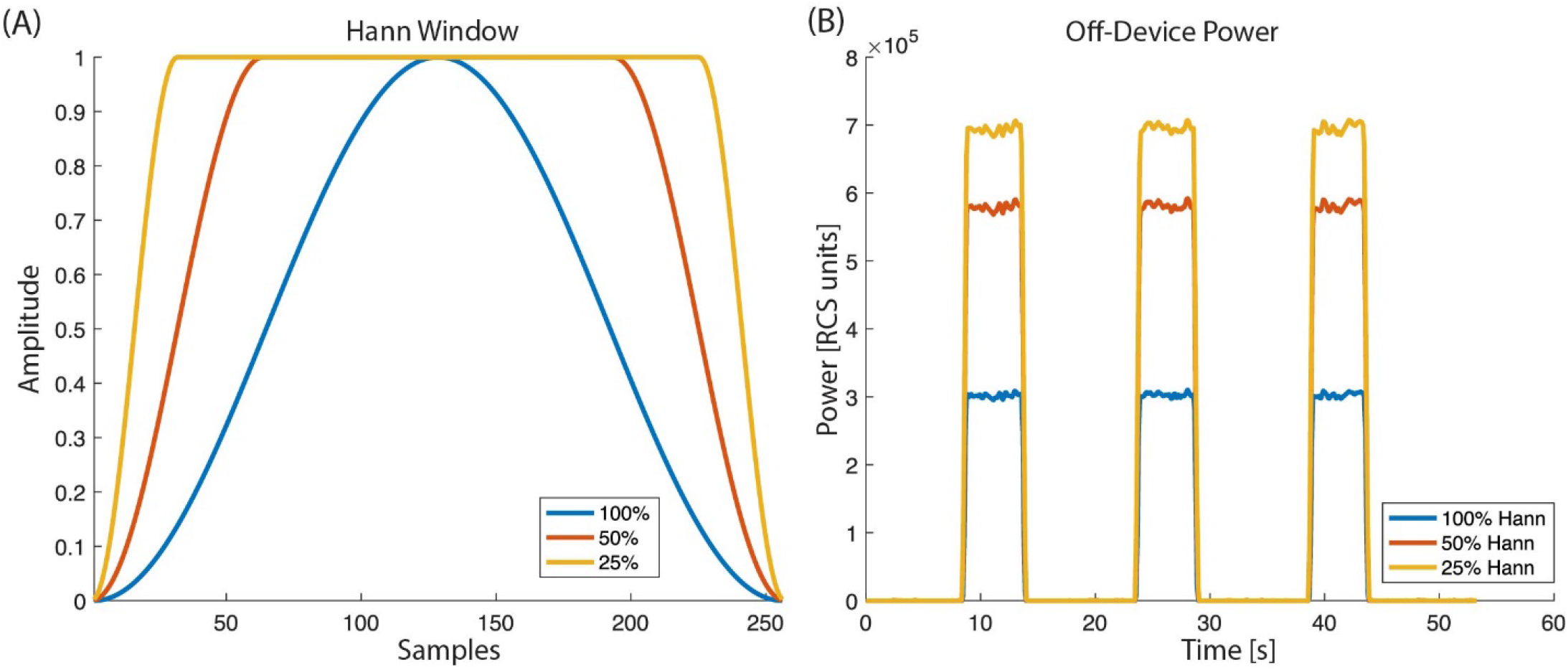
Hann window with ‘window load’ parameter of 25%, 50%, and 100% as selectable by the RC+S FFT power calculation: (A) The shape of the tapper Hann window function. (B) Power calculated off-device based on different Hann window load values.

In Figure 9, the off-device calculation of a benchtop dataset is shown. The time domain raw neural signal s(n) is transformed to the internal on-device units (Eq 1). Then, a window with the size of the FFT is shifted from start to end of the time domain signal using the Hann window (see Eq 2, 3). For each window, the single-sided fast Fourier transform is calculated, and the biomarker power band is computed as the sum of the power of all frequency bins within the defined frequency band.

**Figure 9:**
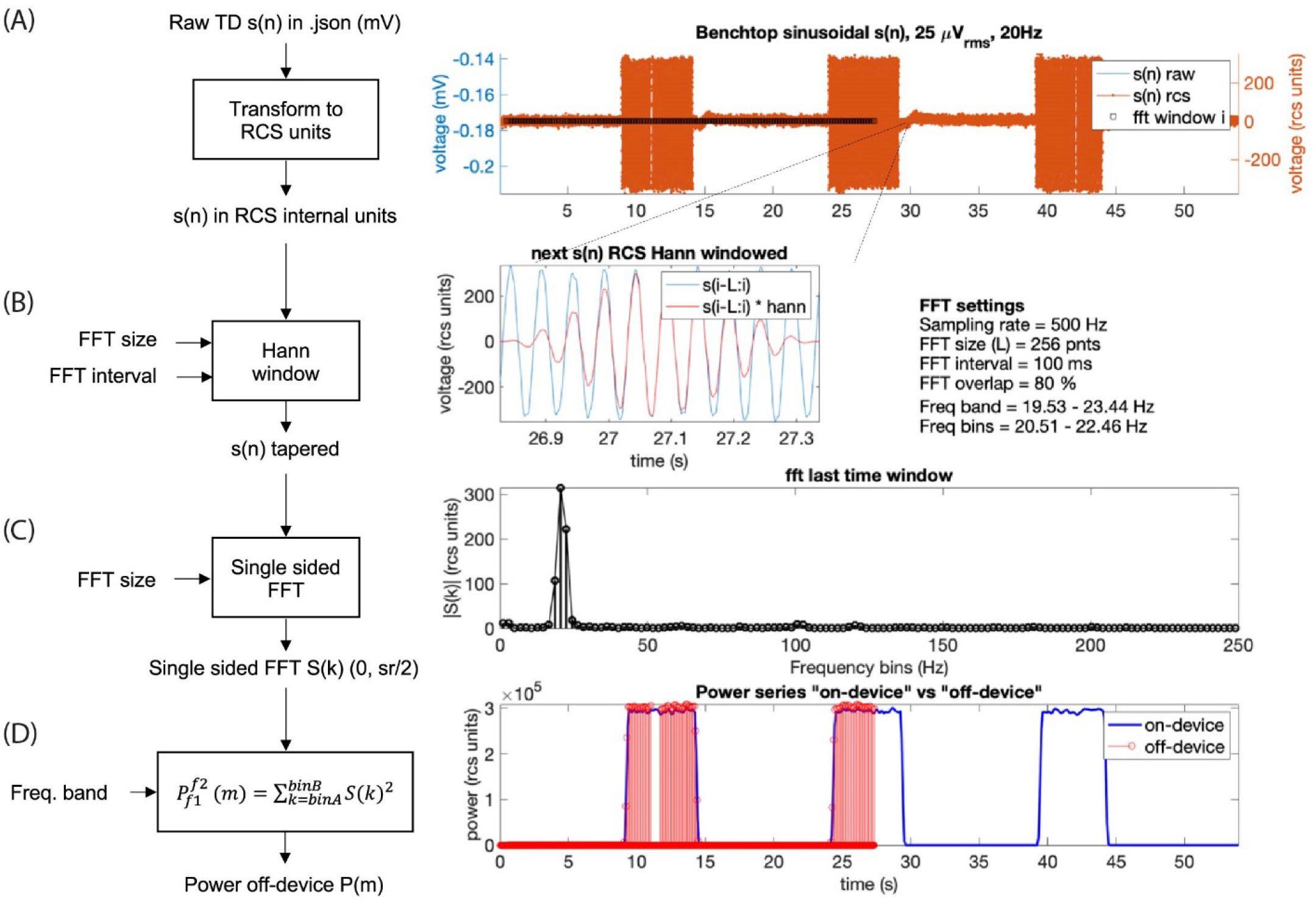
Power calculation off-device replicating the on-device power calculation for a benchtop dataset with three 5 seconds bursts of a 25 microvolts sine wave at 20Hz frequency. The calculation is conducted following 4 steps: (A) The raw time domain neural signal s(n) (mV) is transformed back to internal device units, RCS units (see Eq 1, Table 5). (B) To minimize spectral leakage, a Hann window is applied to each new analysis window of **the** transformed signal s(n). The new analysis window (∼26.9 to 27.3 s) is defined by the size and interval of the FFT (Eq 2, 3). The raw signal within the next time segment is shown in blue and the Hann window tapered signal is in red. (C) A single-sided FFT is applied to the Hann tapered signal s(n) resulting in an amplitude FFT value per each frequency bin of the complete FFT band (0 to ½ sampling rate). For the exact scaling of the single sided FFT see function ‘calculateNewPower’ on https://github.com/openmind-consortium/Analysis-rcs-data. (D) Power is computed as the sum of squares of each FFT amplitude for all frequency bins within the frequency band. The on-device power series is shown in blue and the off-device calculated power, up to the last analyzed window in this graph (∼27s) is depicted in red (using the matlab function stem). The time alignment between the on-device and the off-device signal is accurate as the perfect overlay between sample points at the power signal flanks shows.

In Figure 10, a comparison between the on-device and off-device calculations for a human subject dataset is shown. To assess the difference between the on-device and the off-device calculated power, root mean square error (RMSE) and percentage difference were evaluated, resulting in 284.72 (RCS units) and 1.41% difference, respectively.

**Figure 10:**
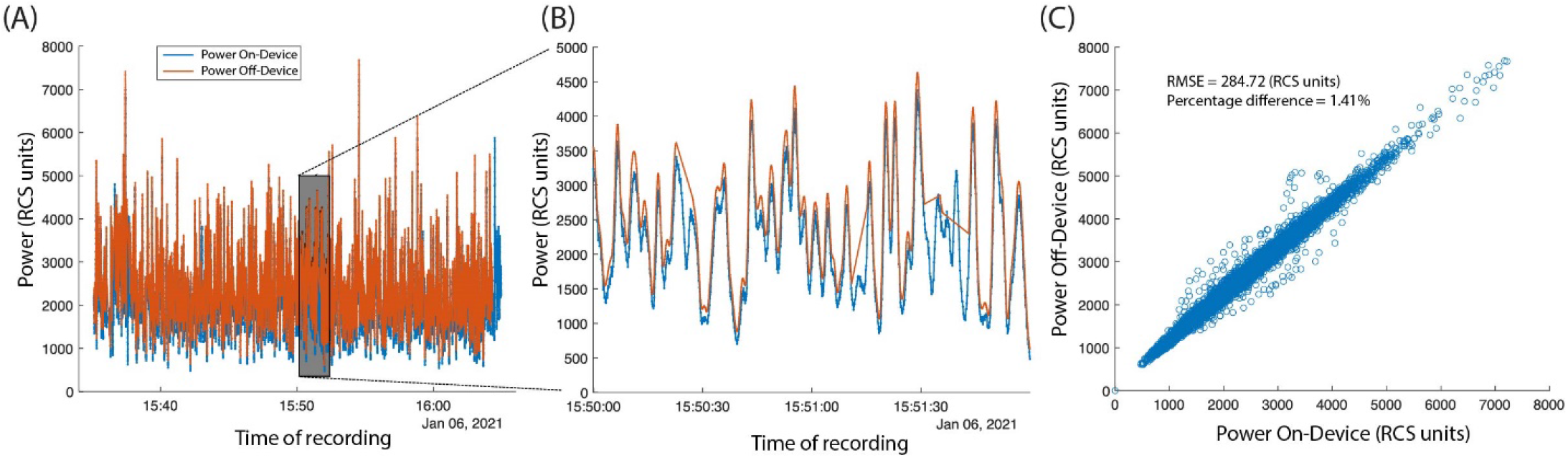
In vivo human data set showing on-device and off-device calculated power series for a given frequency band (8.05 to 12.20 Hz) (A) Overlay of power time series for the ‘on-device’ and the ‘off-device’ calculation. (B) Zoom into a 2-minute segment showing minimal difference between the two power series. (C) The scatter plot showing the fit between the ‘on-device’ and ‘off-device’ power values with RMSE of 284.72 (rcs units) and percentage difference of 1.41%.

## Discussion

DBS is an established or experimental therapy for a number of neurological and psychiatric diseases (Krauss et al. 2021; Mayberg et al. 2005; Lozano et al. 2008; Schlaepfer et al. 2013; Fontaine et al. 2015; Moro et al. 2017; Limousin and Foltynie 2019; Pereira and Aziz 2014; Harmsen et al. 2020; Mallet et al. 2009; Shirvalkar et al. 2020). Originally applied in an open-loop paradigm, there has been a surge of interest in delivering closed-loop or adaptive stimulation in response to disease and symptom biomarkers (Arlotti et al. 2018; Neumann et al. 2014; Hoang and Turner 2019; Provenza et al. 2019). The Medtronic Summit RC+S bidirectional device is being tested in a number of clinical trials for therapeutic stimulation to treat a range of diseases (Table 1). This device is equipped with advanced sense and stimulation capability, including the ability to record multiple data streams simultaneously (e.g. time domain, accelerometry, power, FFT, adaptive detectors), and to stimulate either in open-loop or adaptive mode. The device is powered via a rechargeable battery, thereby allowing patients to stream 24/7 without the need for frequent surgeries to replace a primary cell battery. However, fully leveraging these advanced capabilities is limited if researchers and clinicians cannot efficiently access recorded data in a format amenable to conventional analysis to inform device programming. Here, we provide a toolbox which can ingest raw JSON data from the Medtronic Summit RC+S device and provide key outputs and functionality for users. We import raw time series and metadata from all data streams and decode information to human-readable values. Critically, we compute a common time base such that all data streams can be analyzed together with time alignment. While seemingly simple, the technical specifics of how data packets are transmitted from the INS to the external tablet precluded the ability to easily analyze multiple datastreams together with accurate time alignment prior to this implementation. Prior studies relied on averaging time windows for more coarse alignment or looking at datastreams independently. Because our common time base is in Unix time, it further facilitates the synchronization of Summit RC+S data with external sensors and tasks and event-locked epoching. The ability to analyze all acquired datastreams together is fundamental to both our scientific understanding of neural correlates of disease and for accurately understanding how neural activity, stimulation, and symptoms relate for the clinical management of implanted patients. Similarly, viewing these different datastreams together provides a more comprehensive view of the therapy. Our plotting tool allows for easily customized visualization of one or multiple datastreams.

The Summit RC+S system provides the technological advance to enable embedded adaptive stimulation. Such therapy has been applied in the treatment of epilepsy (Kremen et al. 2018) and Parkinson’s Disease (Swann et al. 2018; Gilron et al. 2021). Across indications, the device is programmed to calculate power within predefined frequency bands, and these values are used to determine the current ‘state’, relative to the predefined detector thresholds. The Summit RC+S has two detectors available when operating in embedded adaptive mode, each with a linear discriminant function that allows for up to 4 input power features. Selecting all the parameters for each computation, detector, and threshold is a challenge in the real-world implementation of this system. Exhaustive testing with patient reports of symptom status (in order to validate performance) is not feasible because of the large parameter space. Therefore, we provide a tool which allows Summit RC+S users to calculate inferred embedded power estimates, off the device, using streamed time domain data. While standard software power calculations can be used to analyze the data for better understanding of neural correlates of symptom status, those computations are less useful in informing programming of the device. Here, we mimic the computation steps performed on the device hardware and firmware in order to obtain values that are comparable to what the device will calculate. The magnitudes of the power values calculated are typically used to set the threshold values in the detector, so it is critical to have off-device computations which do not require a scaling factor or other transform to be comparable to online computations.

Though the Summit RC+S is only accessible via an Investigational Device Exemption with no current plans for commercial release, it has a nine-year life span, and is expected to be implanted in over 130 patients across 7 indications. Given the research volume planned with these patients (estimated to be over $40M in federal funding), a robust toolbox to aid in data analysis and data sharing could prove invaluable for the research groups that will be working on these datasets in the decade(s) to come. New bidirectional sense and stimulation enabled devices continue to enter the market (e.g. Medtronic Percept). We hope the learnings presented via this toolbox can provide guidance to device manufacturers to develop systems which are easily implemented and managed by clinicians and researchers (Borton et al. 2020). While coding of data may be needed to overcome the limited transmission bandwidth available to fully-implanted devices, translation of these codes to human-readable values as early as possible in the user-facing pipeline is desirable. Consistent and streamlined handling of missing data, data streams with different sampling rates, and continuous data with changing parameters are critical for efficient analysis. Thoughtful design at this level will decrease the barrier to entry for new clinicians and researchers, which is common in the medical/academic environment, especially when working with patients who are enrolled in multi-year clinical trials. This is particularly important as more neurological and psychiatric conditions are becoming understood in terms of neurophysiology for both biomarker tracking and adaptive stimulation.

In order to facilitate use and adoption of this toolbox, we provide an extensive README in the shared GitHub repository. We provide example datasets, both patient data (anonymized and shared with informed consent) and benchtop data acquired with known characteristics and input signals, to facilitate user training and to demonstrate features of the toolbox. The repository is actively maintained, with ongoing code review of new features and bug fixes.

The presented toolbox includes three key areas of functionality. Future areas for fruitful development are plentiful. The quantity of raw and processed data from patients implanted with Summit RC+S devices is staggering, and efficient databasing is required. This will facilitate both targeted analyses as well as data mining across patients. The toolbox is currently implemented in Matlab, but in the short term a conversion tool can be written to make the data easily accessible by Python. In the long-term, we seek to implement an open-source data standard for Summit RC+S data, Neurodata Without Borders (NWB). The NWB format provides a documented schema on top of the h5 file format and facilitates data readability, sharing, and archiving. Conversion of raw JSON files from the Summit RC+S directly into NWB was not possible because of the unique packet structure and the need to create a shared timebase across all datastreams. With the functionality of the toolbox presented here, we are now able to begin developing conversion modules to create RC+S NWB files. The power computation module we presented serves as a template for future development of analyses - including a similar off-device implementation of the detector engine which utilizes linear discriminant analysis. Such tools can be applied to data collected prior to chronic implant in order to inform personalized targeting (Allawala et al. 2021).Taken together, we hope this toolbox provides infrastructure on which to continue building shared analysis tools for the ongoing development of stimulation therapy using the Medtronic Summit RC+S for the whole neurophysiology community.

## Acknowledgements

The authors are grateful to members of the OpenMind Consortium, Starr, Chang, and Shirvalkar labs for helpful discussion and testing of the software platform. We thank David Linde and Scott Stanslaski for technical insight into the Summit RC+S device. Devices provided by Medtronic at no cost.

This work was partially funded by UH3NS115631 (KKS, PRS, EFC, PAS), U24NS113637 (RG, PAS), and UH3NS100544 (RG, KHL, SJL, PAS). JA’s salary was paid for by the Swiss National Science Foundation (Early Postdoc Mobility - P2BEP3_188140).

KKS, RG, and JA Conceived of the presented toolbox. KKS, RG, and JA wrote the toolbox code. KKS, RG, JA, KHL, and PRS tested the code toolbox. KKS, RG, and JA wrote the manuscript. PRS, EFC, SJL, and PAS provided data, funding, and mentorship for the project. All authors contributed to manuscript revision, read, and approved the submitted version.

